# A Proteomic Atlas of Senescence-Associated Secretomes for Aging Biomarker Development

**DOI:** 10.1101/604306

**Authors:** Nathan Basisty, Abhijit Kale, Okhee H Jeon, Chisaka Kuehnemann, Therese Payne, Chirag Rao, Anja Holtz, Samah Shah, Vagisha Sharma, Luigi Ferrucci, Judith Campisi, Birgit Schilling

**Affiliations:** The Buck Institute for Research on Aging, Novato, California 94947, USA; Lawrence Berkeley Laboratory, University of California, Berkeley, California 94720, USA; University of Washington, Seattle, Washington 98195, USA; National Institute on Aging, Bethesda, Maryland 20892, USA

## Abstract

The senescence-associated secretory phenotype (SASP) has recently emerged as a driver of, and promising therapeutic target for, multiple age-related conditions, ranging from neurodegeneration to cancer. The complexity of the SASP, typically assessed by a few dozen secreted proteins, has been greatly underestimated, and a small set of factors cannot explain the diverse phenotypes it produces *in vivo*. Here, we present the ‘SASP Atlas’, a comprehensive proteomic database of soluble and exosome SASP factors originating from multiple senescence inducers and cell types. Each profile consists of hundreds of largely distinct proteins, but also includes a subset of proteins elevated in all SASPs. Our analyses identify several candidate biomarkers of cellular senescence that overlap with aging markers in human plasma, including GDF15, STC1 and SERPINs, which significantly correlated with age in plasma from a human cohort, the Baltimore Longitudinal Study of Aging. Our findings will facilitate the identification of proteins characteristic of senescence-associated phenotypes and catalog potential senescence biomarkers to assess the burden, originating stimulus and tissue of origin of senescent cells *in vivo*.

**Abbreviations:** ATV, atazanavir treatment; BLSA, Baltimore Longitudinal Study of Aging; CTL, control; DDA, data-dependent acquisition; DAMP, damage-associated molecular pattern; DIA, data-independent acquisition; eSASP, extracellular vesicle senescence associated secretory phenotype; EVs, extracellular vesicles; IR, X-irradiation; MS, mass spectrometry; RAS, inducible RAS overexpression; SA-β-Gal, senescence-associated β-galactosidase; SEN, senescent; sSASP, soluble senescence associated secretory phenotype.

## Introduction

Cellular senescence is a complex stress response that causes an essentially irreversible arrest of cell proliferation and development of a multi-component senescence-associated secretory phenotype (SASP) [1–4]. The SASP consists of a myriad of cytokines, chemokines, growth factors and proteases that initiate inflammation, wound healing and growth responses in nearby cells [5, 6]. In young healthy tissues, the SASP is typically transient and tends to contribute to the preservation or restoration of tissue homeostasis [5]. However, senescent cells increase with age and a chronic SASP is known or suspected to be a key driver of many pathological hallmarks of aging, including chronic inflammation, tumorigenesis and impaired stem cell renewal [5, 7]. Powerful research tools have emerged to investigate the effect of senescence on aging and disease, including two transgenic p16^Ink4a^ mouse models that allow the selective elimination of senescent cells [8, 9] and compounds that mimic the effect of these transgenes. Data from several laboratories, including our own, strongly support the idea that senescent cells and the SASP drive multiple age-related phenotypes and pathologies, including atherosclerosis [10], osteoarthritis [11], cancer metastasis and cardiac dysfunction [12, 13], myeloid skewing [14, 15], kidney dysfunction [16], and overall decrements in healthspan [17]. Recently, senescent cells were shown to secrete bioactive factors into the blood that alter hemostasis and drive blood clotting [18]. SASP factors therefore hold potential as plasma biomarkers for aging and age-related diseases that are marked by the presence of senescent cells.

To develop robust and specific senescence and aging biomarkers, a comprehensive profile of the context-dependent and heterogeneous SASP is needed. Several types of stress elicit a senescence and SASP response, which in turn can drive multiple phenotypes and pathologies associated with mammalian. These stressors have both shared and distinct secretory components and biological pathways. For example, telomere attrition resulting from repeated cell division (replicative senescence), ionizing radiation, chromatin disruption, and activation of certain oncogenes all can cause senescence-inducing genotoxic stresses, as can genotoxic therapeutic drugs, such as certain anti-cancer chemotherapies [13] and therapies for HIV treatment or prevention [19]. However, while both ionizing radiation and oncogenes lead to DNA double-strand breaks, ionizing radiation uniquely produces clustered oxidative DNA lesions [20] whereas oncogene activation drives DNA hyper-replication and double strand breaks [21]. Whether different senescence-inducers produce similar or distinct SASPs is at present poorly characterized. Thus, a comprehensive characterization of SASP components is critical to understanding how senescent responses can drive diverse pathological phenotypes *in vivo*.

The SASP was originally characterized by antibody arrays, which are necessarily biased, to measure the secretion of a small set of pro-inflammatory cytokines, proteases and protease inhibitors, and growth factors [1,2,4,22]. Subsequently, numerous unbiased gene expression studies performed on different tissues and donors of varying ages suggest that the SASP is more complex and heterogeneous [23], however, a recent meta-analysis of senescent cell transcriptomes confirmed the expression of a few dozen originally characterized SASP factors in multiple senescent cell types [24].

While unbiased transcriptome analyses are valuable, they do not directly assess the presence of *secreted* proteins. Thus, proteomic studies are needed to accurately and quantitatively identify SASP factors as they are present in the secretomes of senescent cells. Recently, a mass spectrometric study reported several SASP factors induced by genotoxic stress [25], but an in-depth, quantitative and comparative assessment of SASPs originating from multiple stimuli and different cell types is lacking. Senescent cells also secrete bioactive exosomes [26, 27] with both protein and miRNA [28] cargos. But, aside from pro-tumorigenic effects [28] and ability to induce paracrine senescence [26, 29], the proteomic content and function of exosomes and small extracellular vesicles (EVs) secreted by senescent cells remains largely unexplored.

In this study, we demonstrate that the SASP is not a single phenotype, but rather is highly complex, dynamic and dependent on the senescence inducer and cell type. Here we also present the “SASP Atlas” (www.SASPAtlas.com), a comprehensive, curated and expanding online database of the soluble secretomes of senescent cells (sSASPs) induced by various stimuli in several cell types. We also present the first comprehensive proteomic analysis of the exosomal SASP (eSASP), which is largely distinct from the sSASP. Our approach leverages an innovative data-independent mass spectrometry workflow to discover new SASP biomarker candidates. The SASP Atlas can help identify candidate biomarkers of aging and diseases driven by senescent cells. We also show that the SASP is enriched for protein markers of human aging and propose a panel of top SASP-based aging and senescence biomarker candidates.

## Results

### Cellular senescence entails extensive changes in the secreted proteome

We established an efficient, streamlined proteomic workflow to discover novel SASP factors. We collected proteins secreted by senescent and quiescent/control primary human lung fibroblasts (IMR90) and renal cortical epithelial cells (Fig 1). Briefly, we induced senescence in the cultured cells by X-irradiation (IR), inducible oncogenic RAS overexpression (RAS), or treatment with the protease inhibitor atazanavir (ATV, used in HIV treatment) and allowed 1-2 weeks for the senescent phenotype to develop, as described [2]. In parallel, control cells were made quiescent by incubation in 0.2% serum for 3 days and were either mock-irradiated or vehicle-treated. Treated and control cells were subsequently cultured in serum-free medium for 24 hours and the conditioned media, containing soluble proteins and exosomes/extracellular vesicles (EVs), was collected. Soluble proteins and exosomes/EVs were separated by ultracentrifugation.

**Fig 1.**
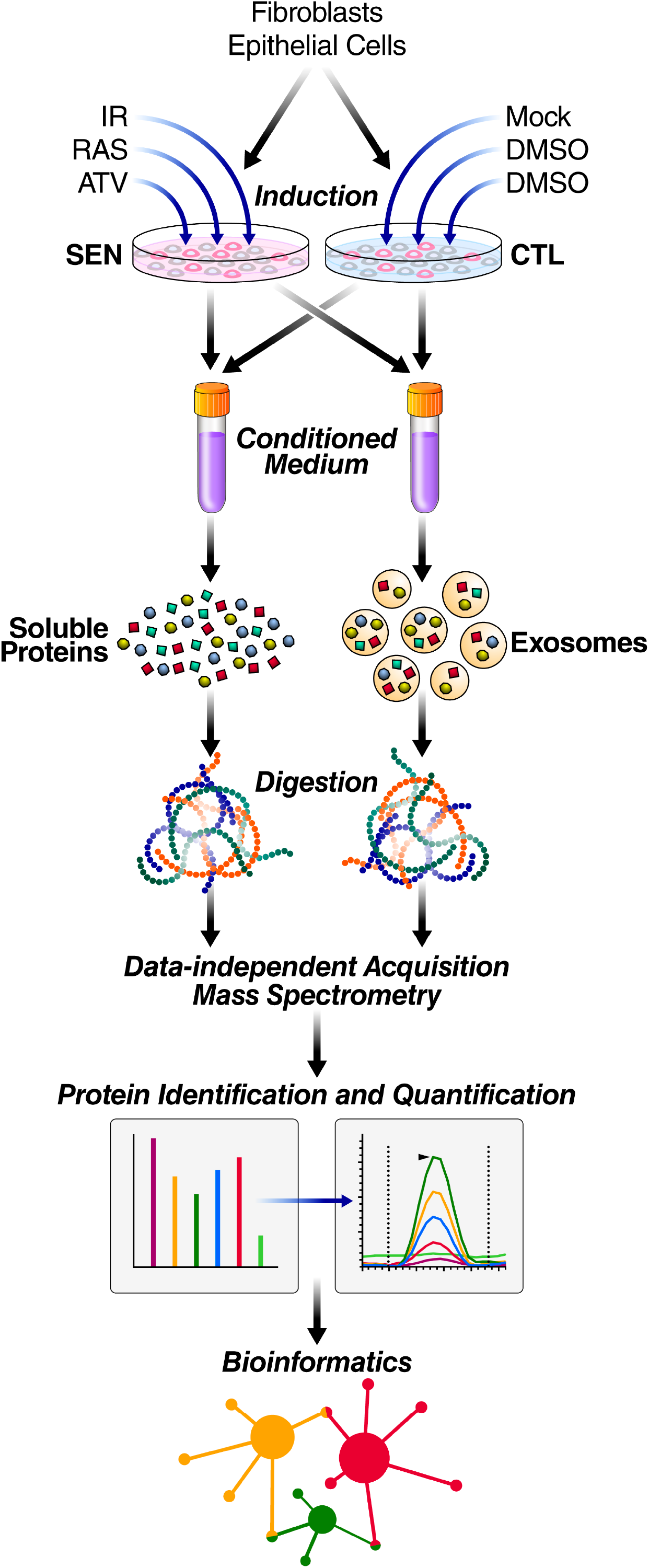
Proteomic workflow for isolation and analysis of secreted proteins and exosomes/EVs. Senescence was induced in cultured primary human lung fibroblasts and renal epithelial cells by either X-irradiation (IR), inducible oncogenic RAS overexpression (RAS), or atazanavir (ATV). Control cells were made quiescent and either mock irradiated or vehicle treated. Soluble proteins and exosomes/EVs were then isolated from conditioned media. Samples were digested and subjected to mass spectrometric analysis (DIA), followed by protein identification and quantification using Spectronaut Pulsar [32] and by bioinformatic, pathway and network analyses in R and Cytoscape [33, 34]. SEN = Senescent, CTL = Control.

The label-free data-independent acquisition (DIA) approach enabled sensitive and accurate quantification of SASP proteins by integrating the MS2 fragment ion chromatograms [30, 31]. We quantitatively compared proteins secreted by senescent cells with controls, and significantly changed proteins (q-value <0.05) that had a fold change of at least 1.5-fold (SEN/CTL) were identified. Proteins secreted at significantly higher levels by senescent relative to quiescent cells were defined as SASP factors. In fact, most proteins were secreted at much higher levels by senescent cells compared to non-senescent cells (Fig 2). Each treatment and control group contained 4-10 biological replicates (see Methods for replicate details and experimental design). Relative protein quantification and statistical details are presented in **Table S1**. Induction of senescence was verified by senescence-associated β-galactosidase (SA-β-Gal) activity and p16INK4a and IL-6 mRNA levels (Fig S1A-C), as described [2]. There was no detectable cell death, as measured by a Sytox Green viability dye assay (Fig S2). X-irradiation and RAS overexpression induced senescence in >90% of cells and ATV induced senescence in about 65% of cells (Fig S1A-B).

**Fig 2.**
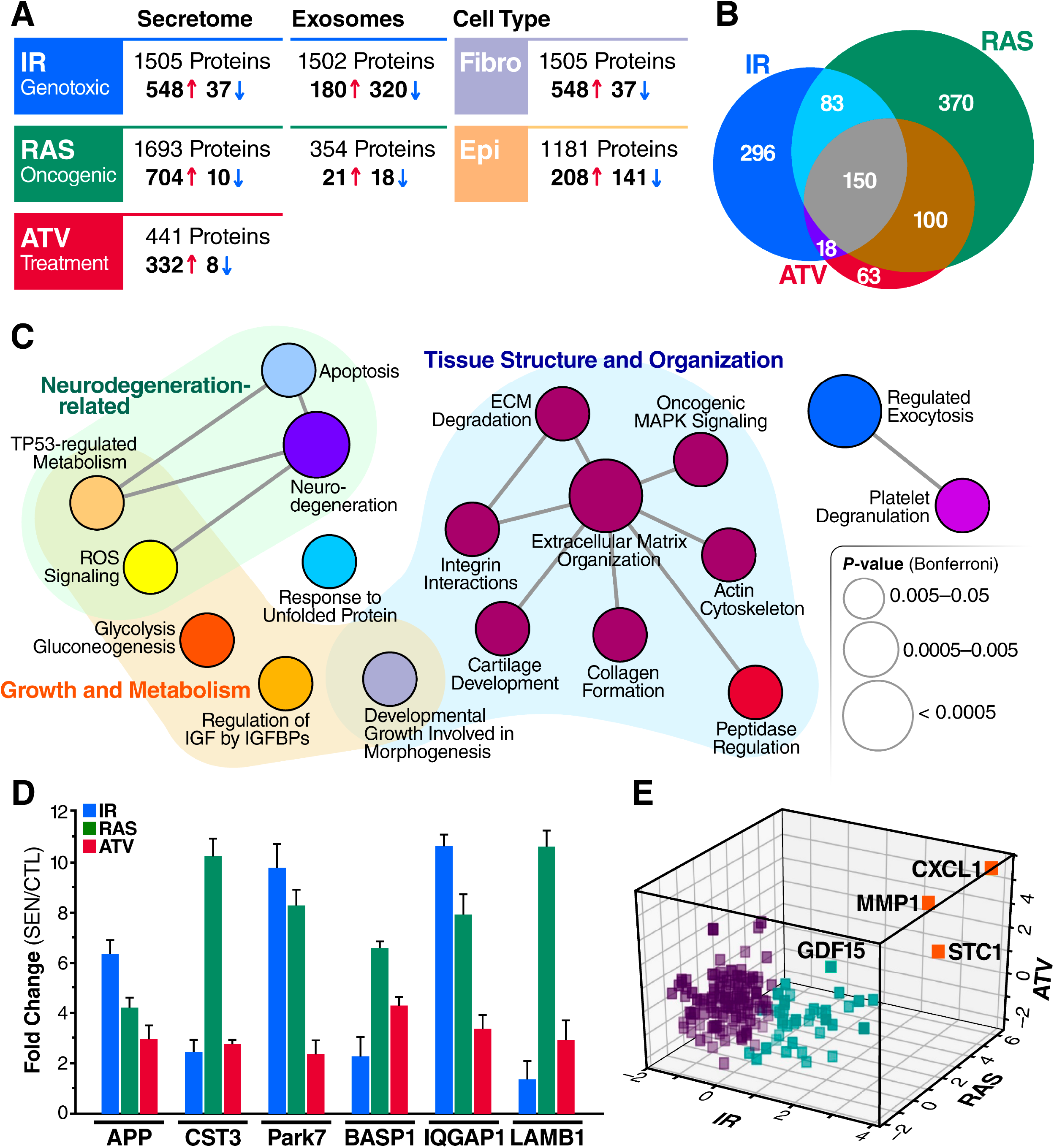
Core sSASP proteins, networks and pathways. (A) Summary of proteins with significantly altered (q-value<0.05) secretion by senescent compared to quiescent cells following genotoxic, oncogenic, or ATV treatment stress in senescent human lung fibroblasts and renal epithelial cells. B) Venn diagram of proteins showing significantly increased secretion in senescent versus non-senescent cells following induction of senescence by IR, RAS or ATV. (C) ClueGO [33] pathway enrichment and network analyses of overlapping sSASPs resulting from each senescence inducer. Pathways of the same color have >= 50% similarity. Connecting lines represent Kappa connectivity scores >40%. (D) Secretion levels of proteins in the neurodegeneration pathway, expressed as log2 fold change of senescent versus control cells. (E) Unsupervised K-means clustering of proteins significantly increased in the sSASPs of all inducers based on the magnitude of the protein changes (log2-change) in senescent versus control groups and partitioned into three clusters. IR = X-irradiation, RAS = RAS oncogene overexpression, ATV = atazanavir treatment.

This unbiased proteomic profiling identified up to ∼1700 secreted proteins, a large fraction of which were up- or down-regulated following induction of senescence by IR, RAS or ATV (Fig 2). Between 340 to 714 proteins changed significantly in response to each inducer. As expected, most of the significantly changed proteins were markedly upregulated in senescent, compared to quiescent, cells, but, interestingly, a minority were downregulated (Fig 2A). Notably, the protein cargo of exosomes/EVs released by senescent cells was distinct compared to that from non-senescent cells (Fig 2A), indicating the existence of an exosome/EV SASP (eSASP) in addition to the sSASP.

Most changes in the sSASP, independent of inducer, exhibited increased secretion by senescent cells, with only 1-6% of proteins secreted at lower levels. In contrast, one-half to two-thirds of all significant protein changes in exosomes/EVs from senescent fibroblasts were decreased relative to quiescent cells (Fig 2A). For renal epithelial cells, the sSASP comprised a more even mix of proteins with significantly lower or higher relative secretion. The magnitude of the fold-changes in the sSASP were generally higher in fibroblasts than in renal epithelial cells, regardless of inducer (Fig 2A). For example, 531 of significant protein changes in the fibroblast sSASP were >2-fold, compared to 138 in the renal epithelial cell sSASP. However, for renal epithelial cells an additional 212 proteins showed significant changes between 1.5- and 2-fold increase or decrease.

For each cell type and fraction, we also measured the secretion of known SASP factors (Fig S1D). These factors included chemokines (CXCLs), high mobility group box 1 protein (HMGB1), IGF binding proteins (IGFBPs), matrix metalloproteinases (MMPs), lamin B1 (LMNB1), and tissue inhibitors of metallopeptidase (TIMPs). In fibroblasts, all previously identified SASP factors were elevated, regardless of the senescence inducer. However, while expression of p16INK4a, IL-6 and SA-β-Gal were also elevated in renal epithelial cells (Fig S1A-C), most classical SASP proteins were either decreased or unchanged, except for IGFBP2. This finding suggests that fibroblast SASP markers do not necessarily pertain to other cell types. Similarly, within exosomes/EVs secreted by senescent fibroblasts, nearly all previously identified key SASP factors were either absent, unchanged or decreased, and none were consistently elevated in response to more than one inducer (Fig S1D).

### Senescence-inducing stimuli drive largely distinct secretory phenotypes

To determine how different senescence-inducing stimuli affect the SASPs, we compared the sSASP from human primary fibroblasts induced to senesce by IR, RAS or ATV. Strikingly, the sSASP was largely distinct among inducers, with an overlap of 150 proteins among 1091 total increased proteins and no overlap among decreased proteins (**Table S2**). Thus, most SASP protein components and corresponding changes were highly heterogenous and not shared among inducers (Fig 2B).

To determine whether there are core pathways associated with the SASPs, we performed pathway and network analyses on overlapping proteins in the sSASPs of each inducer (Fig 2C). The largest pathway associated with all inducers related to tissue and cell structure, including extracellular matrix, cytoskeleton, integrins and peptidase activity. Interestingly, neurodegeneration and three related pathways with high agreement (kappa score >40%) – apoptosis, ROS signaling and TP53-regulated metabolism - were also enriched among the overlapping sSASP proteins (Fig 2C-D). Among the neurodegeneration proteins were amyloid precursor protein (APP) and cystatin 3 (CST3), related to Alzheimer’s pathogenesis and risk [35, 36], as well as Parkinsonism-associated deglycase (PARK7 or DJ1) [37] (Fig 2D). This enrichment of neurodegeneration-associated proteins and pathways suggests that senescent cells contribute to neurodegenerative diseases, for which these sSASP factors might serve as biomarkers, regardless of the senescence-inducing stimuli.

To distill the overlapping ‘core’ sSASP proteins into primary components, we performed an unsupervised machine learning analysis (Fig 2E). K-means clustering analysis uncovered three primary clusters among core sSASP components. Strikingly, one cluster, consisting of just three proteins – chemokine C-X-C motif ligand 1 (CXCL1), matrix metallopeptidase 1 (MMP1), and stanniocalcin 1 (STC1) – were highly represented in the sSASPs of all inducers, suggesting these proteins might serve as surrogate markers of the sSASP. Of note, STC1, among the top sSASP proteins, is a previously unidentified SASP factor and a secreted hormone with many disease associations [38–43]. Our analyses also validate MMP1 and CXCL1 as SASP markers.

### sSASP is largely distinct in composition and regulation in fibroblasts and epithelial cells

We compared the secretomes of lung fibroblasts and renal epithelial cells to determine the cell-type specificity of the sSASP. The sSASP of these cells were largely distinct (Fig 3A-B). Among the proteins increased in the sSASP of each cell type, 9-23% overlapped, and the magnitude of the changes by renal epithelial cells were, in most cases, lower than in fibroblasts regardless of the senescence inducer, although it is possible that senescent fibroblasts secrete more protein overall than epithelial cells in response to stress. Interestingly, 20-30% of proteins significantly decreased in the sSASP of renal epithelial cells overlapped with proteins significantly increased in the fibroblast sSASP (Fig 4B). Among the epithelial factors that changed oppositely to the fibroblast factors were IGFBPs, TIMPs 1 and 2, CXCL1 and most SERPINs (Fig 2C). In all, 17 sSASP factors were shared between all senescence inducers and cell types we examined (**Table S3**).

**Fig 3.**
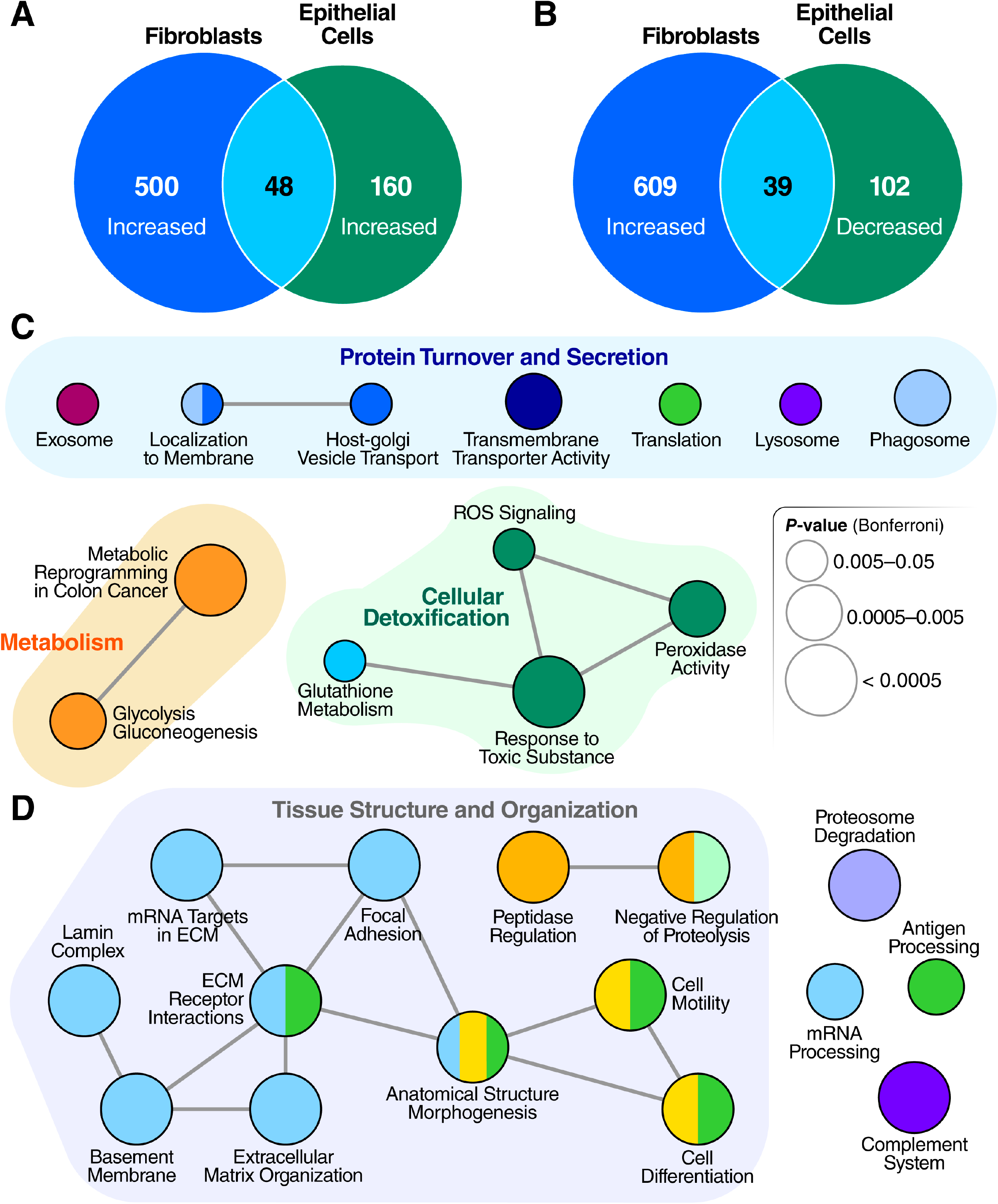
Epithelial cells and fibroblasts have distinct sSASPs. (A) Venn diagram comparing proteins significantly increased in the sSASPs of senescent fibroblasts and epithelial cells, both induced by X-irradiation (q< 0.05). (B) Venn diagram comparing protein increases in the fibroblast sSASP vs *decreases* in the epithelial sSASP. (C) Pathway and network analysis of secreted proteins significantly increased by senescent fibroblasts and epithelial cells. (D) Pathway and network analysis of proteins significantly increased in the fibroblast sSASP but significantly decreased in the epithelial cell sSASP.

**Fig 4.**
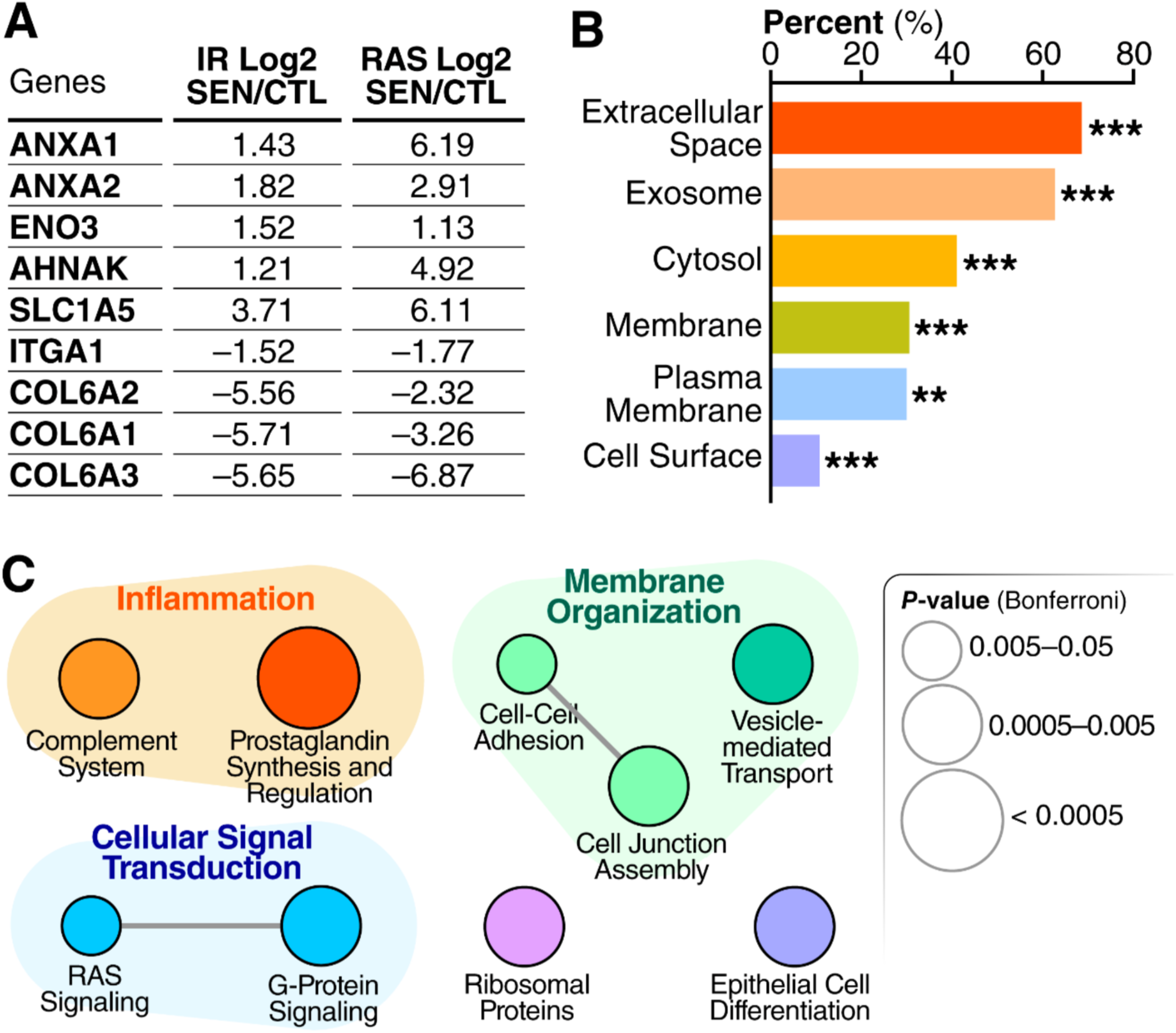
Cellular senescence alters exosome/EV features and composition. (A) Table showing overlapping significant protein changes in exosomes/EVs secreted by senescent cells induced by IR vs RAS (q < 0.05). (B) Enrichment analysis of gene-ontology/cellular compartments overrepresented among protein contents of exosomes/EVs released by senescent cells. (C) Network analysis of pathways and functions unique to the eSASP.

Pathway and network analysis of proteins increased in the sSASPs of fibroblasts and epithelial cells (Fig 3C) showed that most pathways belonged to one of three general categories: protein turnover and secretion, primary metabolism, and cellular detoxification. While not as apparent on a molecule-by-molecule basis, many pathways were commonly enriched in both the epithelial and fibroblast sSASPs (Fig 3C and Fig 2C), including vesicle-mediated transport and exosomes, glycolytic metabolism, and cellular detoxification. Of notable exceptions, pathways enriched uniquely by epithelial cells included protein translation and degradation (lysosome and phagosome).

Surprisingly, most renal epithelial sSASP proteins with significantly *lower* secretion by senescent cells were enriched in pathways related to tissue and cell structure, adhesion and motility (Fig 3D). This finding contrasts with previous reports and our own analyses of fibroblasts (Fig 2C), in which these pathways were increased, regardless of inducer. The irradiated epithelial sSASP also had significantly lower levels of proteins involved in RNA processing, in contrast to increased RNA metabolism in the irradiated fibroblast sSASP. Additionally, the epithelial sSASP was significantly depleted in proteins related to proteasome degradation, antigen processing and the complement system.

Damage-associated molecular patterns (DAMPs, also known as alarmins or danger signals) are released from cells in response to internal and external stress, and are components of the SASP [44]. HMGB1 is a founding member of the DAMPs, a prominent SASP marker, and, along with calreticulin (CALR), an important driver of inflammation [44]. Our analysis identified increased secretion of multiple DAMPs, including HMGB1 and CALR, by senescent fibroblasts under all senescence inducers (Table 1). However, the secretion of DAMPs was unchanged or significantly reduced by senescent epithelial cells, demonstrating that some defining SASP components vary depending on cell type.

**Table 1:** DAMPs are a core component of the fibroblast sSASP.

|  | **Log2(SEN/CTL)** |  |  |  |
| --- | --- | --- | --- | --- |
|  | **<u>IR (Fibroblasts)</u>** | **<u>RAS</u>** | **<u>ATV</u>** | **<u>IR (Epithelial)</u>** |
| **HMGB1** | 2.47 | 0.59 | 2.46 | NS |
| **CALR** | 1.22 | 0.51 | 1.32 | -1.00 |
| **CD44** | 2.25 | 1.20 | 1.92 | -0.69 |
| **S100A11** | 0.56 | 1.35 | 1.88 | NS |
| **LGALS3BP** | 1.46 | 1.76 | 1.79 | -1.19 |
| **VCAN** | 1.80 | 1.32 | 0.98 | -1.42 |
| **TNC** | 1.64 | 1.40 | 2.46 | 0.29 |
| **HSPA5** | 2.03 | 3.93 | 1.78 | -0.34 |
| **HSP90AB1** | 5.01 | 2.69 | 1.65 | NS |
| **HSPA8** | 2.49 | 2.98 | 1.46 | 0.32 |
| **HSPA1A** | 2.96 | 2.4 | 1.45 | 0.54 |
| **HSP90AA1** | 4.94 | 3.42 | 1.34 | NS |
| **HSP90B1** | 2.67 | 1.61 | 0.66 | -0.27 |

All changes are significant ( $q < 0.05$ ) unless denoted NS (Not Significant). SEN = Senescent, CTL = Quiescent Control, IR = X-irradiation, RAS = oncogenic RAS overexpression, ATV = Atazanavir treatment.

### Exosome/EV proteomic signatures are altered by cellular senescence

Because proteins are also secreted as extracellular vesicle cargo, we hypothesized that senescent cells would show significant changes in this fraction, which we term the exosome/EV SASP (eSASP). We used ultracentrifugation to enrich conditioned media for exosomes and small EVs released by quiescent and senescent fibroblasts induced by X-irradiation and oncogenic RAS overexpression (Fig 1). We confirmed the quality of exosome/EV purified fractions by measuring presence of multiple EV-specific markers, including CD63 and CD9 [45], and by particle counting and size distribution analysis (Fig S4). Exosomes/EVs from senescent fibroblasts showed a strikingly altered protein composition compared with exosomes from non-senescent cells (Fig 2A).

To determine whether the characteristics of exosomes/EVs from senescent cells are altered, we analyzed particle number and size distribution of exosomes/EVs secreted into the culture medium of senescent and non-senescent cells over a 24-hour period. On average, senescent cells released a greater number of vesicles -- about 68 per cell compared to 49 per control cell (Fig S4B). However, the mean diameter, size distribution of senescent and control exosomes/EVs were similar (Fig S4B-C). Further work using senolytics may validate whether the number, size, and other characteristics of secreted exosome/EVs are indicators of senescent cell burdens in humans.

The protein content of exosomes/EVs released by IR-vs RAS-induced senescent fibroblasts was largely distinct, sharing only 9 significantly altered proteins (Fig 4A). Exosomes/EVs were reported to contain protein signatures of their originating cells [28, 46], offering a unique opportunity to identify senescence biomarkers with a degree of cell type specificity. Thus, exosome/EV proteins might distinguish senescent cells of different origins or resulting from different stressors. The membranes of exosomes are also representative of the originating cells [28, 46]. Indeed, about 30% of all the exosome/EV proteins that increased upon senescence are plasma membrane proteins (Fig 4B), suggesting that exosomes/EVs might also identify cell type origins through their cell-surface proteins. In addition to enrichment of proteins involved in membrane organization, such as cell adhesion and cell junction assembly proteins, the eSASP is uniquely enriched with signaling pathways not found in the sSASP, such as RAS signaling, G-protein signaling, and prostaglandin synthesis and regulation (Fig 4C). Full lists of proteins secreted by senescent exosomes are in **Table S1**.

### The SASP contains potential aging and disease biomarkers

As a driver of many aging and disease phenotypes, the SASP could include known biomarkers of aging and age-related diseases. A recent biomarker study identified 217 proteins that are significantly associated with age in human plasma (adjusted p<0.00005) [47]. Of these, 20 proteins (9.2%) were present in the originally-defined SASP [2]. Strikingly, multiple newly identified SASP factors from our present study were also identified in the study of human plasma [47] (Fig 5). Of all the originally-defined SASP factors and unique SASP proteins that we identify here, 101 proteins were also identified as markers of aging in human plasma (46.5% of all plasma aging markers) (Fig 5A, C-D, **Table S4**). Considering the originally defined SASP in addition our newly identified “core SASP” (SASP components resulting from all senescence inducers), the number of age-associated plasma proteins that are also SASP proteins is 40, or 18.4% of plasma aging markers (Fig 5B-D, **Table S4**). Thus, plasma biomarkers of aging are highly enriched with SASP factors.

**Fig 5.**
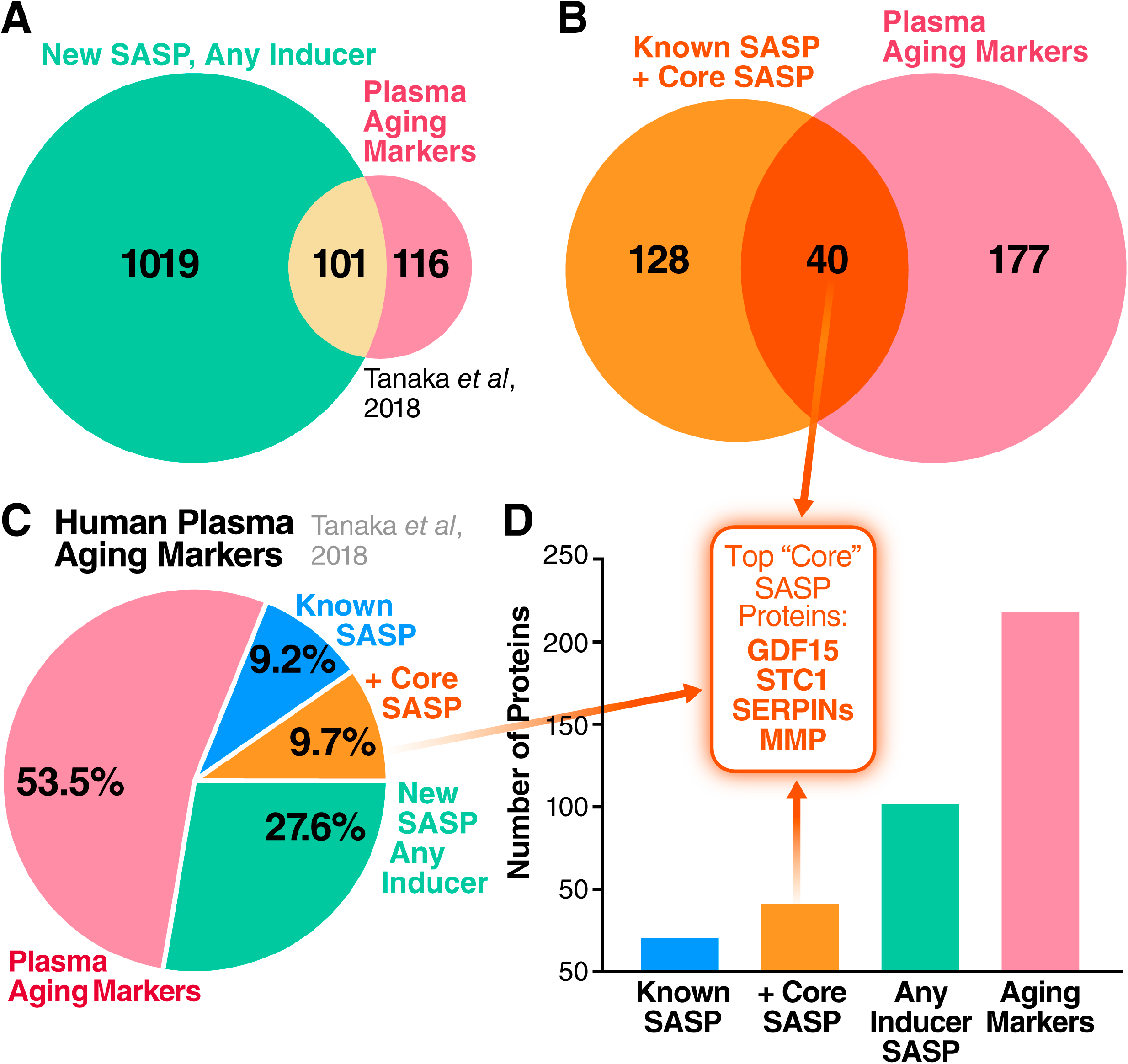
Human plasma aging markers are enriched for SASP proteins. (A) Venn diagram comparing SASP factors secreted by at least one of IR-, RAS-, or ATV-induced senescent cells with markers of aging identified in human plasma [47]. (B) Overlap between the core SASP (proteins secreted following all senescence-inducing stimuli) and plasma aging markers. (C) Pie chart showing the proportion of known SASP factors, newly identified core SASP factors, and SASP factors found among plasma markers of aging in humans. (D) Number of proteins contained in the originally identified SASP, core SASP, non-core SASP, and markers of aging in human plasma [47] (p<0.00005). Top core SASP factors GDF15, STC1, SERPINs, and MMP1 are among the plasma aging markers.

Complement and coagulation cascade proteins [18], particularly protease inhibitors such as SERPINs, were also noted as prominent plasma biomarkers of aging [47]. These proteins and their pathway networks were robustly altered in the SASPs of cells induced to senesce by all the tested stressors (Fig 2C, Fig 6A). The protein having the strongest association with aging [47], GDF15 (r=0.82), was among the most highly secreted proteins in the sSASP induced by IR, RAS and ATV in fibroblasts, and in epithelial cells induced by IR (Fig 6D). Increased secretion of top core SASP biomarkers SERPINE1, MMP1, STC1, and GDF15 was confirmed by western blotting in RAS-induced senescent cells compared to controls (Fig S3). The enrichment of aging and disease biomarkers in the secretomes of senescent cells supports their link to a wide spectrum of age-related diseases.

**Fig 6.**
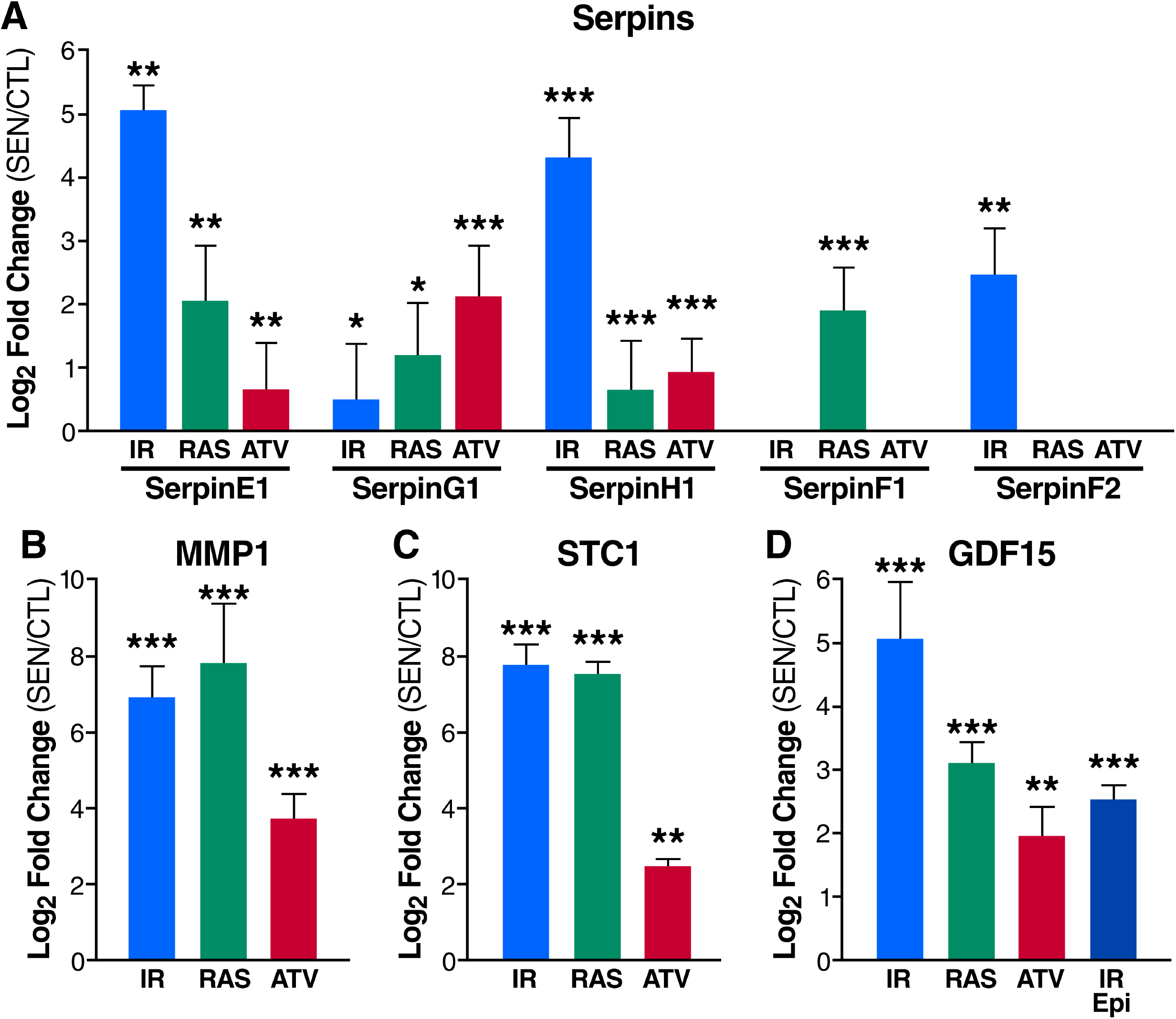
The SASP contains aging and disease biomarkers. (A) Serpins are secreted at high levels by senescent fibroblasts induced by IR, RAS or ATV. (B) MMP1 and (C) STC1 are among the most highly secreted proteins by senescent fibroblasts. (D) The plasma aging biomarker GDF15 is increased in the sSASPs of fibroblasts induced to senesce by IR, RAS and ATV and epithelial cells induced to by IR. IR = X-irradiation, RAS = RAS oncogene overexpression, ATV = atazanavir treatment, Epi = renal epithelial cells. *q < 0.05, **q < 0.01, ***q < 0.001.

## Discussion

Here we present SASP Atlas (www.SASPAtlas.com), the first proteome-based database of SASPs. This database contains the contents of exosome/EV and soluble secretomes, in addition to SASPs originating from multiple senescence-inducing stresses and two distinct cell types. The SASP Atlas will be continuously updated with SASP profiles from new cell types and senescence, including paracrine (or bystander) senescence [48, 49], as well as temporal dynamics of the SASP – all generated by our laboratories.

Our proteomic analysis leverages a modern data-independent acquisition (DIA or SWATH) mass spectrometry workflow, which comprehensively acquires label-free, quantitative peptide (MS1) and fragment-level (MS2) data for all peptides in each sample [30–32,50,51]. DIA workflows are not limited by the stochastic peptide MS/MS sampling biases characteristic of traditional data-dependent acquisition (DDA) mass spectrometry. In addition to the SASP Atlas database, we provide panels of SASP factors on Panorama Web, a freely-available web repository for targeted mass spectrometry assays [52, 53]. These resources can be used as a reference and guide to identify and quantify SASP factors that may be associated with specific diseases, and to develop aging and disease-related biomarkers (Fig 7).

**Fig 7.**
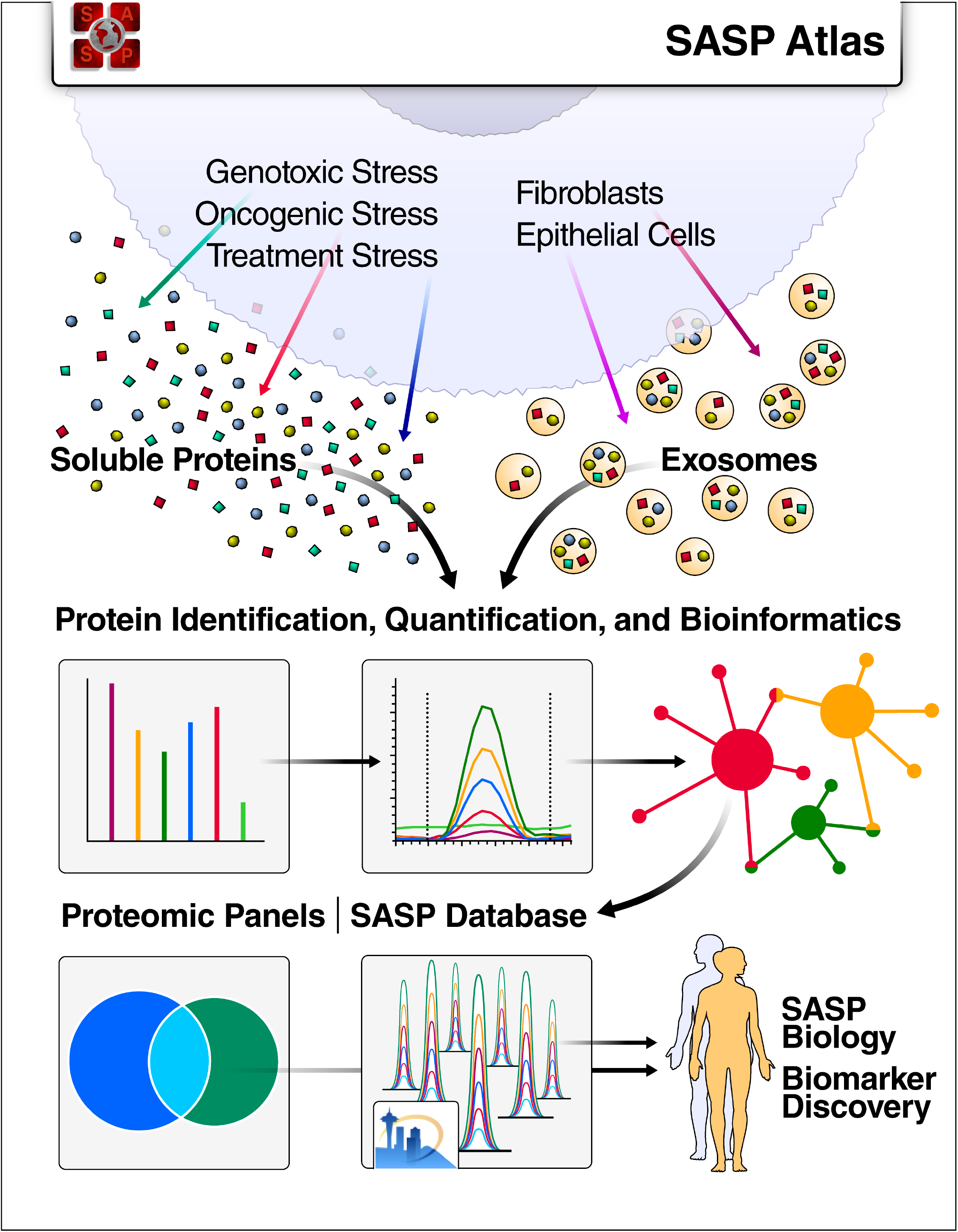
SASP Atlas: A Comprehensive Resource for Senescence-Associated Secretory Phenotypes. SASP Atlas (www.SASPAtlas.com) is a curated and freely-available database of the secretomes of senescent cells, including both the soluble and exosome SASP, that can be used to identify SASP components or biomarker candidates for senescence burden, aging and related diseases.

SASP profiles are needed to develop senescence biomarkers in human plasma or other biofluids, and for identifying individuals to treat with, and measuring the efficacy of, senescence-targeted therapies such as senolytics. Translating senescence- and SASP-targeted interventions to humans will require a comprehensive profile of SASPs, both to identify their deleterious components and to develop human biomarkers to assess senescent cell burden. The SASP, as originally identified, comprised ∼50 cytokines, chemokines, growth factors, and proteases that were detected by biased methods (e.g., antibody arrays) and/or transcriptional analyses [1–4,24]. While these comprehensive analyses are valuable in describing the overall phenotype of senescent cells, proteomic analyses are complimentary in both confirming transcriptional changes and identifying and quantifying novel SASP factors that are not apparent at the mRNA level. For example, a recent meta-analysis of senescent cell transcriptomes [24] identified >1,000 genes with increased expression specifically in senescent cells induced by IR or oncogenic RAS, and >700 ‘core’ senescence genes (increased expression following all senescence inducers tested). Our analysis identified 548, 644, and 143 proteins in the IR, RAS and ‘core’ SASP, respectively, that were previously unreported at the RNA level (Fig S5). We expect that the number and nature of these SASP core proteins will change as we and others interrogate additional cell types and senescence inducers, and we will continue to curate the interactive SASP Atlas. Additionally, the secretion of SASP factors, such as HMGB1 and other DAMPs, is not generally transcriptionally driven.

DAMP receptor-bearing cells, including cells of the innate immune system, recognize extracellular DAMPs as signals to promote inflammatory and fibrotic responses. Increased circulating DAMPs are hypothesized to play a role in aging [54, 55], particularly the age-related inflammation termed ‘inflammaging’ [56]. DAMPs can also serve as biomarkers of a number of diseases, including trauma and cardiovascular, metabolic, neurodegenerative, malignant and infectious diseases [54,57,58]. In addition, our top “core sSASP” biomarker candidates, have been identified as disease biomarkers in human studies. For example, human cohort studies have recently reported GDF15 as a biomarker of cardiovascular disease, cardiovascular and cancer mortality and morbidity, renal disease, and all-cause mortality independent of cardiovascular mortality [59–65]. Additionally, two of the top “core sSASP” proteins identified by an unbiased k-means clustering algorithm – STC1 and MMP1 (Fig 5C-D) – were reported as significant aging biomarkers [47]. In addition to aging, MMP1 has been identified as a biomarker for several cancers, pulmonary fibrosis and potentially Alzheimer’s disease [66–69], whereas STC1 has been identified as a diagnostic and prognostic biomarker for cancers, pulmonary fibrosis, renal ischemia/reperfusion injury and Alzheimer’s disease [38–43].

Our quantitative unbiased *proteomic* analysis of senescent fibroblasts and epithelial cells reveals a much larger and diverse SASP than initially reported. These SASP profiles contribute a number of new potential senescence, aging and disease biomarkers. In addition to general senescence biomarkers, many proteins will likely be specific to cell-type and originating stimulus. Thus, biomarkers present in human patients *in vivo* will likely vary depending on the affected tissue, originating cell types, and senescence stimuli. Therefore, comprehensive quantitative profiles of the SASP under a variety of physiological conditions will provide biomarker candidates with a higher degree of selectivity to specific pathologies in humans.

## Materials and Methods

### Reagents and Resources

A full list of reagents and resources, including vendors and catalog numbers, are available in a Reagent and Resource Table (**Table S5**). Further information and requests for resources and reagents should be directed to the Lead Contact, Birgit Schilling.

### Human Cell Culture and Primary Cell Lines

IMR-90 primary human lung fibroblasts (ATCC #CCL-186) were cultured in Dulbecco’s Modified Eagle’s Medium (DMEM, Gibco #12430-054) supplemented with penicillin and streptomycin (5000 U/mL and 5000 μg/mL, Gibco #15070063) and 10% Fetal Bovine Serum (FBS, Gibco #2614079). Primary human renal epithelial cells (ATCC PCS400011) were cultured in Renal Epithelial Cell Basal Medium (Female, ATCC #PCS-400-030). Both cell types were maintained at 37°C, 10% CO_2_ and 3% O_2_.

### Induction of Senescence

#### X-irradiation

Senescence was induced by ionizing radiation (IR;10 Gy X-ray). Quiescent control cells were mock irradiated. Senescent cells were cultured for 10 days to allow development of the senescent phenotype, and quiescent cells were cultured in 0.2% serum for 3 days. Cells were then washed with PBS (Gibco #10010-023) and placed in serum- and phenol red-free DMEM (Gibco #21063-029) and conditioned media was collected after 24 hours.

#### RAS overexpression

RAS^v12^ was cloned in pLVX vector (Lenti-X™ Tet-On from Clontech #632162) to make inducible lentiviruses, which were used to infect early passage IMR-90 cells (PD-30). Transduced cells were selected in puromycin (1 μg/ml) for 24 hours. For induction of RAS^v12^, cells were treated with 1 μg/ml doxycycline in DMSO (Sigma # D9891) for 4 (early time point) or 7 days. Doxycycline was replaced after every 48 hours. Subsequently, cells were washed with PBS and placed in serum- and phenol red-free DMEM and conditioned media was collected after 24 hours.

#### Atazanivir treatment

Cells were cultured in appropriate media containing 20 µM Atazanavir, which is a clinically relevant dose, or vehicle (DMSO) for 9 (early timepoint) or 14 days. Subsequently, cells were washed with PBS and placed in serum- and phenol red-free DMEM and conditioned media was collected after 24 hours.

### Isolation of Secreted Soluble Proteins and Exosomes/EVs

Proteins secreted into serum-free medium over a 24-hr period were collected. An ultracentrifugation protocol was used to separate the exosome and small extracellular vesicle fraction from the soluble protein fraction [70]. Briefly, conditioned medium was centrifuged at 10,000 x g at 4° C for 30 minutes to remove debris. The supernatant was then centrifuged at 20,000 x g at 4° C for 70 minutes to remove microvesicles followed by ultracentrifugation at 100,000 x g at 4° C for 70 minutes to pellet exosomes. The exosome-depleted supernatant was saved as the sSASP. The exosome pellet was then washed twice with PBS and ultracentrifuged again at 100,000 x g at 4° C for 70 minutes before resuspending in PBS and saved as the eSASP.

### Proteomic Sample Preparation

#### Chemicals

Acetonitrile (#AH015) and water (#AH365) were from Burdick & Jackson. Iodoacetamide (IAA, #I1149), dithiothreitol (DTT, #D9779), formic acid (FA, #94318-50ML-F), and triethylammonium bicarbonate buffer 1.0 M, pH 8.5 (#T7408) were from Sigma Aldrich, urea (#29700) was from Thermo Scientific, sequencing grade trypsin (#V5113) was from Promega and HLB Oasis SPE cartridges (#186003908) were from Waters.

#### Protein concentration and quantification

Samples were concentrated using Amicon Ultra-15 Centrifugal Filter Units with a 3 kDa cutoff (MilliporeSigma #UFC900324) as per the manufacturer instructions and transferred into 8M urea/50 mM triethylammonium bicarbonate buffer at pH 8. Protein quantitation was performed using a BCA Protein Assay Kit (Pierce #23225).

#### Digestion

Aliquots of each sample containing 25-100 µg protein were brought to equal volumes with 50 mM triethylammonium bicarbonate buffer at pH 8. The mixtures were reduced with 20 mM DTT (37°C for 1 hour), then alkylated with 40 mM iodoacetamide (30 minutes at RT in the dark). Samples were diluted 10-fold with 50 mM triethylammonium bicarbonate buffer at pH 8 and incubated overnight at 37°C with sequencing grade trypsin (Promega) at a 1:50 enzyme:substrate ratio (wt/wt).

#### Desalting

Peptide supernatants were collected and desalted with Oasis HLB 30 mg Sorbent Cartridges (Waters #186003908, Milford, MA), concentrated, and re-suspended in a solution containing mass spectrometric ‘Hyper Reaction Monitoring’ retention time peptide standards (HRM, Biognosys #Kit-3003) and 0.2% formic acid in water.

### Mass Spectrometry Analysis

Samples were analyzed by reverse-phase HPLC-ESI-MS/MS using the Eksigent Ultra Plus nano-LC 2D HPLC system (Dublin, CA) combined with a cHiPLC system directly connected to an orthogonal quadrupole time-of-flight SCIEX TripleTOF 6600 or a TripleTOF 5600 mass spectrometer (SCIEX, Redwood City, CA). Typically, mass resolution in precursor scans was ∼ 45,000 (TripleTOF 6600), while fragment ion resolution was ∼15,000 in ‘high sensitivity’ product ion scan mode. After injection, peptide mixtures were transferred onto a C18 pre-column chip (200 µm x 6 mm ChromXP C18-CL chip, 3 µm, 300 Å, SCIEX) and washed at 2 µl/min for 10 min with the loading solvent (H_2_O/0.1% formic acid) for desalting. Peptides were transferred to the 75 µm x 15 cm ChromXP C18-CL chip, 3 µm, 300 Å, (SCIEX), and eluted at 300 nL/min with a 3 h gradient using aqueous and acetonitrile solvent buffers.

All samples were analyzed by data-independent acquisitions (DIA), specifically using variable window DIA acquisitions [71]. In these DIA acquisitions, windows of variable width (5 to 90 m/z) are passed in incremental steps over the full mass range (m/z 400-1250). The cycle time of 3.2 sec includes a 250 msec precursor ion scan followed by 45 msec accumulation time for each of the 64 DIA segments. The variable windows were determined according to the complexity of the typical MS1 ion current observed within a certain m/z range using a SCIEX ‘variable window calculator’ algorithm (more narrow windows were chosen in ‘busy’ m/z ranges, wide windows in m/z ranges with few eluting precursor ions) [31]. DIA tandem mass spectra produce complex MS/MS spectra, which are a composite of all the analytes within each selected Q1 m/z window. All collected data was processed in Spectronaut using a pan-human library that provides quantitative DIA assays for ∼10,000 human proteins [72].

### Cell Viability Assays

Cell viability was assessed with SYTOX Green Nucleic Acid Stain (Invitrogen #S7020). Senescent and control cells were incubated for 24 hours in serum-free medium containing SYTOX Green with continuous imaging. Cell death was quantified by counting total SYTOX Green positive nuclei during the 24-hour time-lapse video.

### Senescence-Associated β-Galactosidase Staining

Senescence-associated beta-galactosidase (SA-β-gal) activity was determined using the BioVision Senescence Detection Kit (Cat# K320-250). For each experiment, approximately 100–150 cells were counted.

### RNA Extraction and Quantitative Real-Time PCR

Total RNA was prepared using the PureLink Micro-to-Midi total RNA Purification System (Invitrogen # 12183018A), according to the manufacturer’s protocol. Samples were first treated with DNase I Amp Grade (Invitrogen #18068015) to eliminate genomic DNA contamination. RNA was reverse transcribed into cDNA using a High-Capacity cDNA Reverse Transcription Kit (Applied Biosystems #4368813), according to the manufacturer’s protocol. Quantitative RT-PCR (qRT-PCR) reactions were performed as described using the Universal Probe Library system (Roche). Actin and tubulin predeveloped TaqMan assays (Applied Biosystems) were used to control for cDNA quantity. qRT-PCR assays were performed on the LightCycler 480 System (Roche).

The primers and probes were as follows:

Human actin F 5’-CCAACCGCGAGAAGATGA; R 5’-TCCATCACGATGCCAGTG, UPL probe #64

Human tubulin F 5’-CTTCGTCTCCGCCATCAG; R 5’-TTGCCAATCTGGACACCA, UPL Probe #58

Human IL-6 F 5’-GCCCAGCTATGAACTCCTTCT; R 5’-GAAGGCAGCAGGCAACAC, UPL Probe #45

Human p16^INK4a^ F 5’-GAGCAGCATGGAGCCTTC; R 5’-CGTAACTATTCGGTGCGTTG, UPL Probe #34

### Exosome Characterization and Size Distribution Analysis

Protein determination is performed on exosomes/EVs isolated by ultracentrifugation by direct absorbance and 20 µg of protein is used for input for the MacsPlex Exosome Kit (Miltenyi) assay. These exosomes are enriched for CD63, CD9, and CD81 surface proteins using antibody beads. This pool of exosomes is then probed for 34 other surface markers used for analysis and comparison across samples. Particle diameter and concentration were assessed by tunable resistive pulse sensing (TRPS) on an IZON qNano Nanoparticle Characterization instrument using a NP150 nanopore membrane at a 47 calibration with 110 nm carboxylated polystyrene beads at a concentration of 1.2×10^13 particles/mL (Zen-bio, Inc.).

### Processing, Quantification, and Statistical Analysis of MS Data

DIA acquisitions were quantitatively processed using the proprietary Spectronaut v12 (12.020491.3.1543) software [32] from Biognosys. A pan-human spectral library was used for Spectronaut processing of the DIA data [72]. Quantitative DIA MS2 data analysis was based on extracted ion chromatograms (XICs) of 6-10 of the most abundant fragment ions in the identified spectra. Relative quantification was performed comparing different conditions (senescent versus control) to assess fold changes. The number of replicates for each experiment are as follows: X-irradiated fibroblasts, 4 senescent and 4 control replicates; X-irradiated epithelial cells, 5 senescent and 5 control replicates; 4 day RAS-induction fibroblasts, 10 senescent and 10 control replicates; 7 day RAS-induced fibroblasts, 6 senescent and 6 control replicates; atazanavir-treated fibroblasts, 3 senescent (9 days treatment), 3 senescent (14 days treatment), and 4 control replicates; X-irradiated fibroblast exosomes, 5 senescent and 5 control replicates; 7 day RAS-induced fibroblast exosomes, 6 senescent and 6 control replicates. Significance was assessed using FDR corrected q-values<0.05.

### Pathway and Network Analysis

Gene ontology, pathway, and network analysis was performed using the GlueGO package, version 2.5.3, in Cytoscape, version 3.7.1 [33, 34]. Curated pathways for enrichment analysis were referenced from the following databases: GO Biological Function, GO Cellular Compartment, Kegg pathways, WikiPathways, and Reactome Pathways. For gene ontology data, testing was restricted to pathways with experimental evidence (EXP, IDA, IPI, IMP, IGI, IEP). The statistical cutoff for enriched pathways was Bonferroni-adjusted p-values < 0.01 by right-sided hypergeometric testing. Pathway-connecting edges were drawn for kappa scores > 40%. Kappa scores are a measure of inter-pathway agreement among observed proteins that indicate whether pathway agreement is greater than expected by chance based on shared proteins. Pathways with the same color indicate >= 50% similarity in terms.

### K-Means Clustering

Unsupervised clustering was performed in Python with Scikit-learn, a module integrating a wide range of machine learning algorithms [73]. Datasets were pre-processed with the StandardScaler function and clustered with the KMeans algorithm.

### Data Visualization

Heatmaps were visualized in R using the heatmap.2 function in the ‘gplots’ package [74]. Venn diagrams were constructed using the “VennDiagram” package [75]. Color palettes in R were generated with the “RColorBrewer” package [76]. Pathway and network visualizations were generated and modified using the GlueGO package in Cytoscape [33, 34].

## Data availability

All raw files are uploaded to the Center for Computational Mass Spectrometry, MassIVE and the ProteomeXchange Consortium and can be downloaded using the following link ftp://massive.ucsd.edu/MSV000083750 (MassIVE ID number: MSV000083750, ProteomeXchange ID number: PXD013721). Data uploads include the protein identification and quantification details, spectral library and FASTA file used for mass spectrometric analysis. SASP proteomic profiles are available on Panorama (https://panoramaweb.org/project/Schilling/SASP_Atlas_Buck/begin.view?), a repository for targeted mass spectrometry assays generated in Skyline software [52, 53]. All data are available for viewing and downloading on SASP Atlas (www.saspatlas.com).

## MassIVE

MSV000083750 (ftp://massive.ucsd.edu/MSV000083750)

## ProteomeXchange

PXD013721 (http://proteomecentral.proteomexchange.org/cgi/GetDataset?ID=PXD013721)

## SASP Atlas

www.SASPAtlas.com

## SASP Panels

https://panoramaweb.org/project/Schilling/SASP_Atlas_Buck/begin.view?

## Acknowledgments

We thank John C.W. Carroll for graphical support generating figures.

## Author Contributions

Conceptualization, N.B., B.S., J.C. L.F.; Cell Culture, N.B., A.K., O.J., C.K., T.P.; RNA expression and Activity Assays, N.B., A.K., O.J., C.K. T.P.; Proteomic Sample Preparation, N.B., T.P., A.H., S.S.; Data Analysis, N.B., B.S., A.H., S.S.; Pathway and Network Analysis, N.B., C.R.; Visualization, N.B., C.R.; Web Database, V.S., C.R., N.B, B.S.; Writing and Editing, N.B., B.S., J.C., L.F.; Funding Acquisition, B.S., J.C.

## Supporting Information: 5 figures, 5 tables

**Fig S1:**
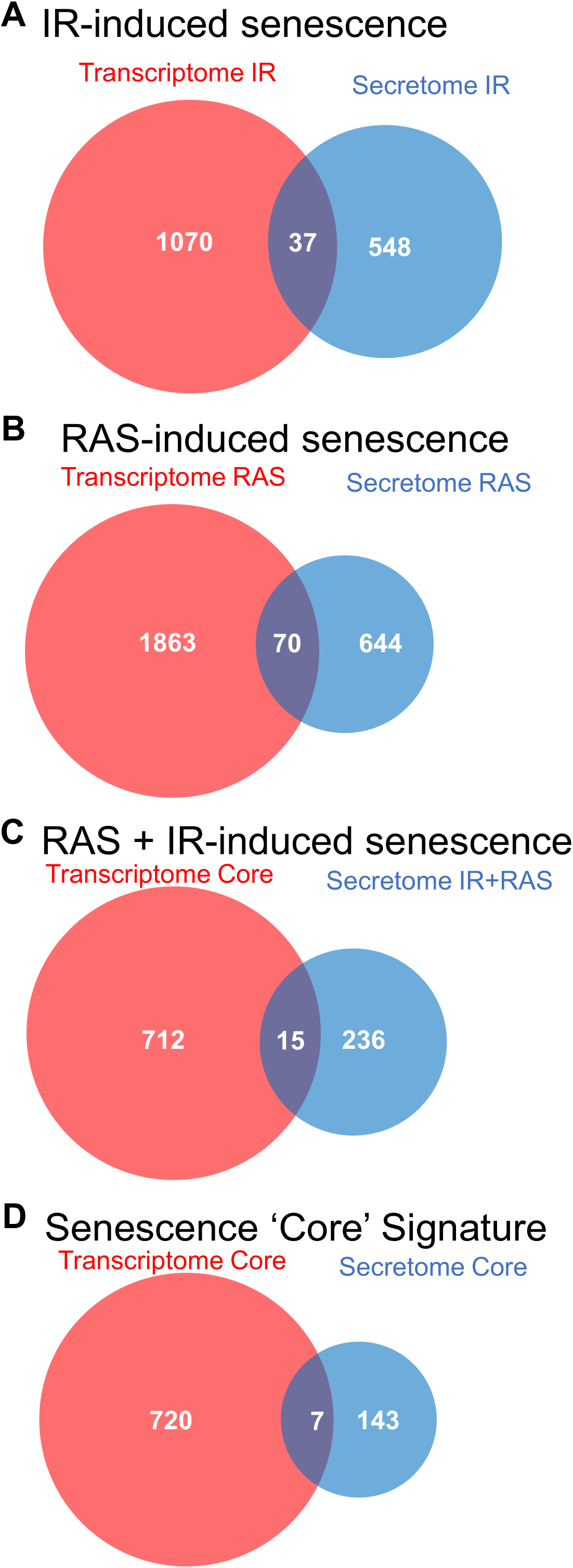
Senescence markers induced by IR, RAS and ATV. (A) Representative images of SA-β-Gal staining of senescent and control (quiescent) primary human lung fibroblasts and renal epithelial cells following induction of senescence by either IR, RAS or ATV. (B) Quantification of SA-β-Gal positive cells. (C) Levels of p16^INK4a^ and Il-6 mRNAs determined by qPCR and expressed as fold change of senescent over control (red line) cells. (D) Commonly reported SASP factors for each inducer, cell type and fraction. IR = X-irradiation, RAS = RAS oncogene overexpression, ATV = atazanavir treatment, Fib = fibroblasts, Epi = renal epithelial cells.

**Fig S2:**
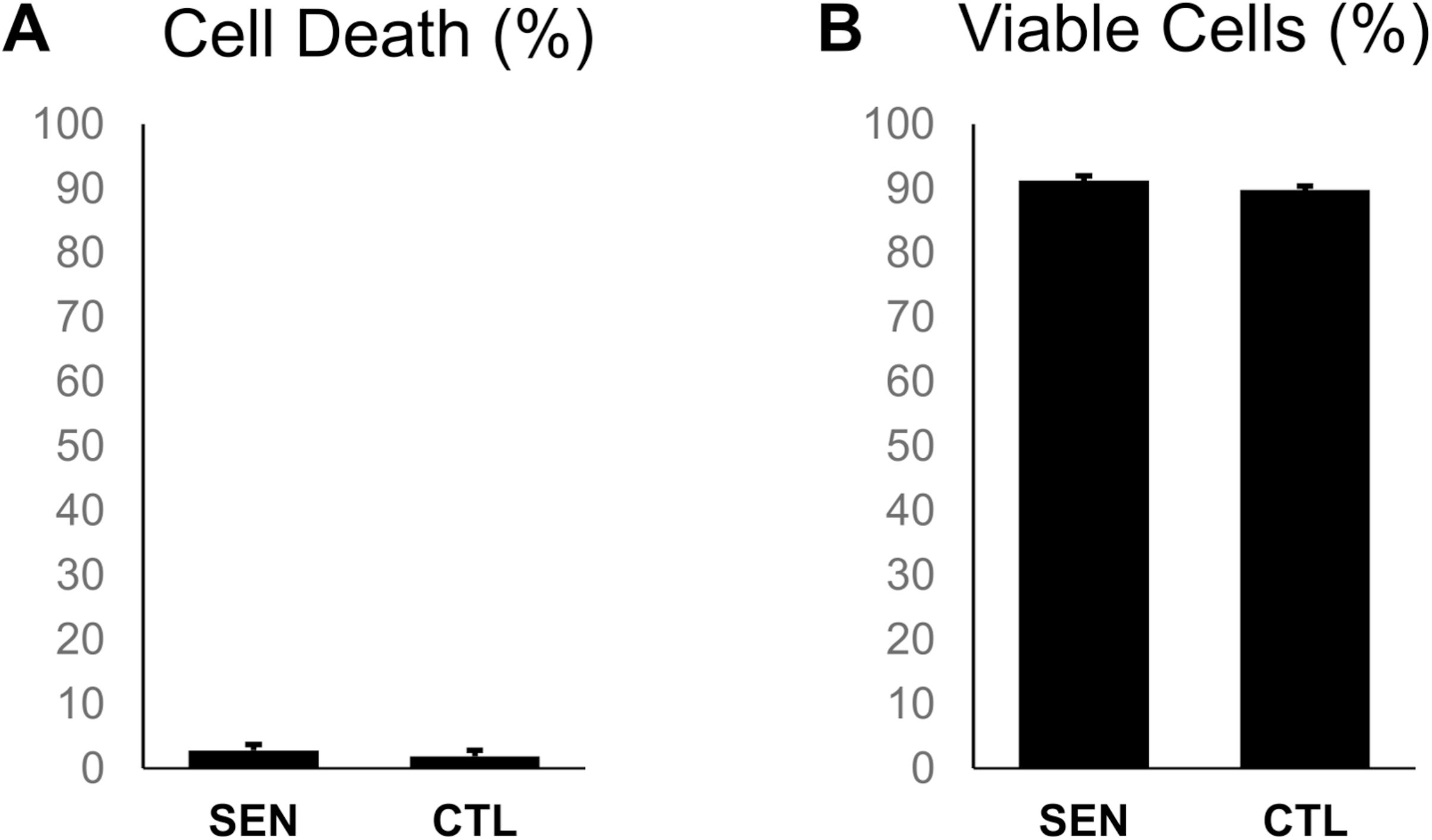
Cell viability assays. (A) Amount of cell death over a 24-hour period as determined by Sytox Green viability dye assay. (B) Fraction of viable cells measured by exclusion of propidium iodide fluorescence, assessed by flow cytometry.

**Fig S3:**
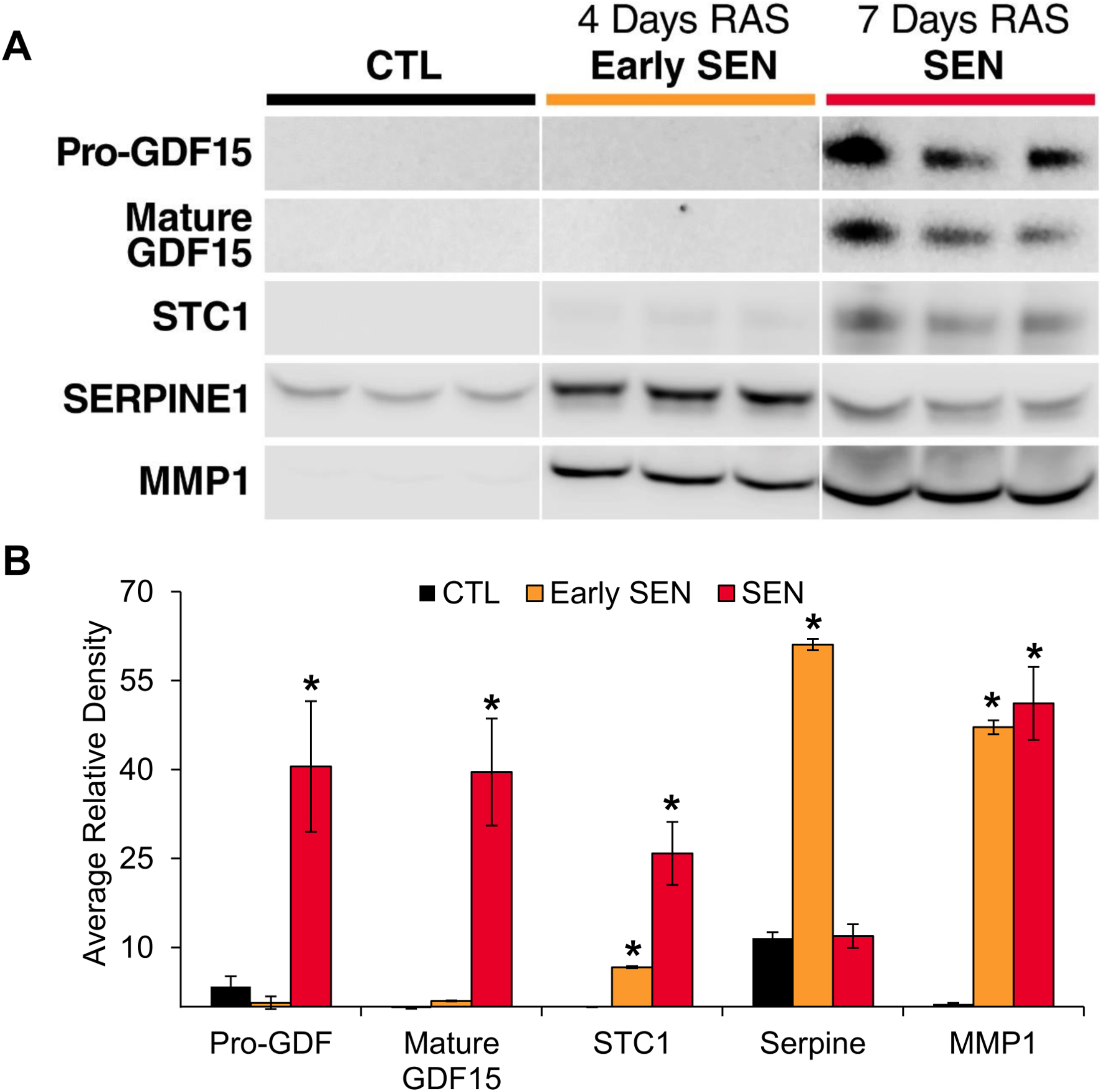
Western Blot Confirmation of top core SASP factors. (A) Western blot exposures of top core SASP factors, GDF15, STC1, SERPINE1, and MMP1, in non-senescent control fibroblasts, early senescent fibroblasts (4 days RAS induction), and fully senescent fibroblasts (7 days RAS induction). (B) Densitometry analysis of western blot. *P-value < 0.05 versus CTL.

**Fig S4:**
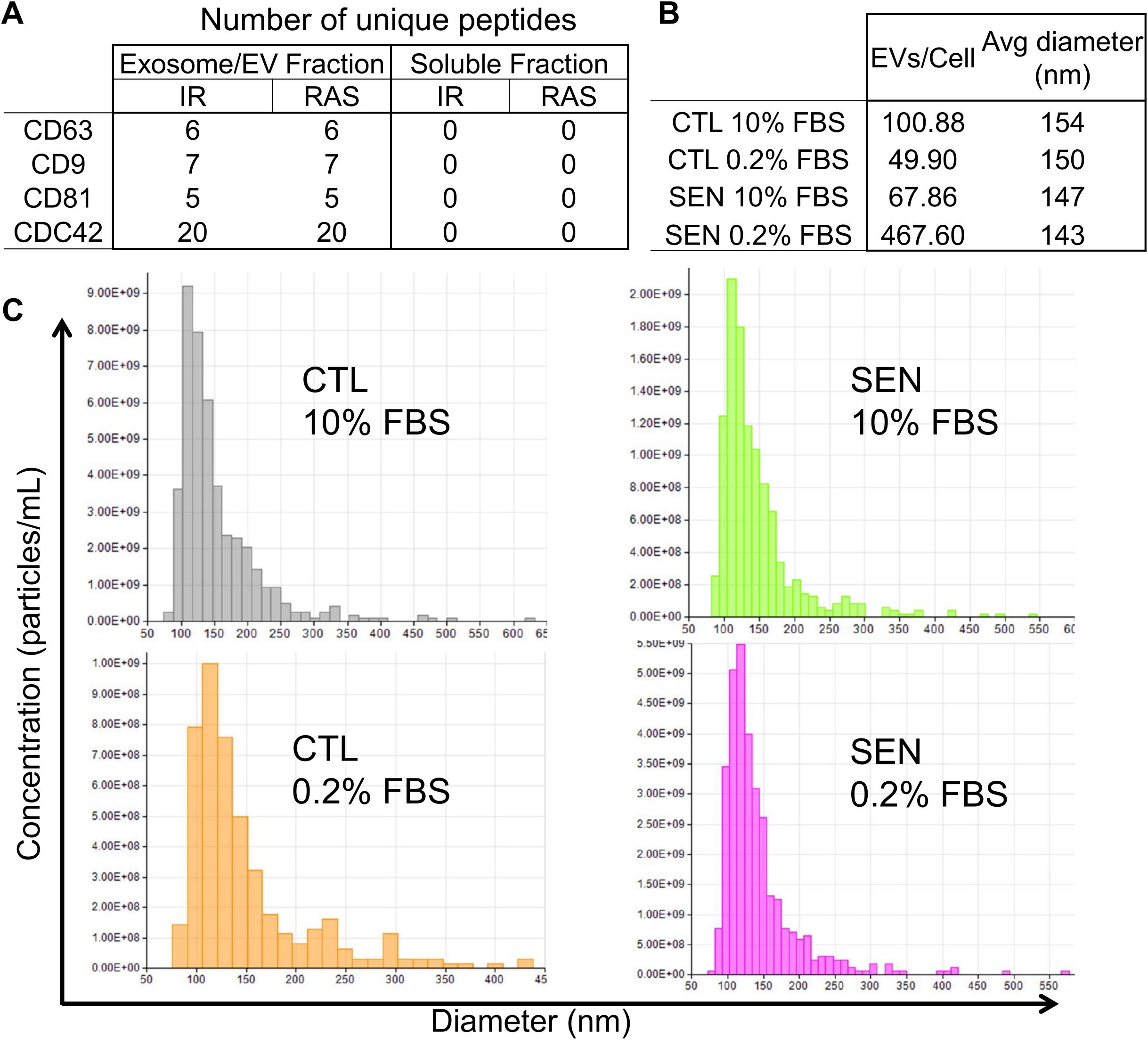
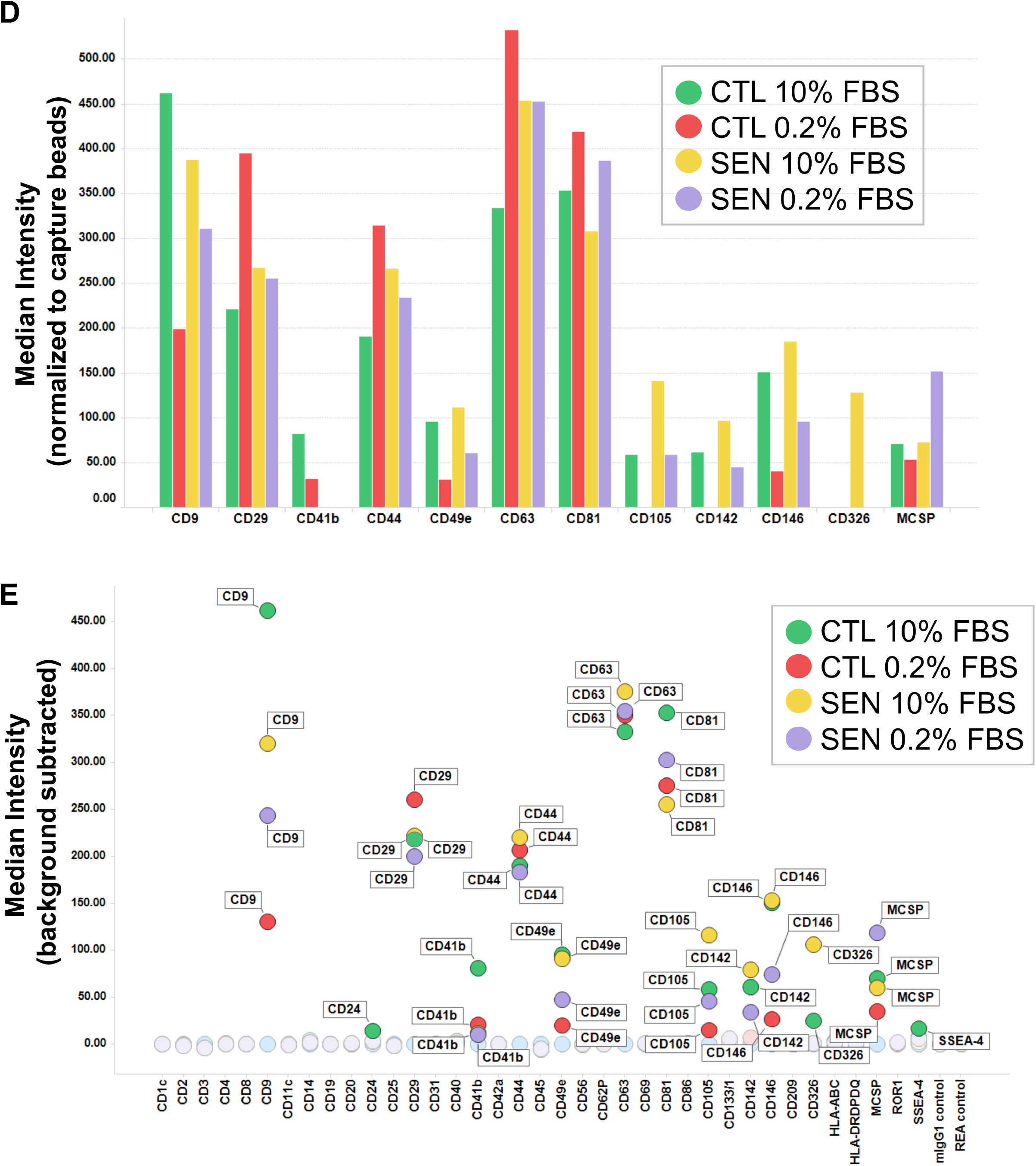
Exosome/EV proteomic markers and size distribution analysis. (A) Table of exosome and EV-specific markers identified in exosome and soluble fractions of fibroblasts by mass spectrometry. Multiple peptides from defining exosome/EV markers were identified in the exosome fractions of RAS and IR-induced senescence experiments but none were detected in the soluble fractions. (B) Table showing EVs secreted per cell and average EV diameter in senescent and control cells in complete (10% FBS) medium and low-serum (0.2% FBS) medium. (C) Size distribution analysis of EVs secreted by senescent and control cells in complete and low-serum medium. (D) Exosome/EV-specific markers detected in isolated EV fractions in each treatment group, as measured by MACSPlex exosome detection kit. (E) Median levels of every surface marker measured in exosome/EV fractions by MACSPlex exosome detection kit. IR = X-irradiation, RAS = RAS oncogene overexpression, FBS = fetal bovine serum.

**Fig S5:**
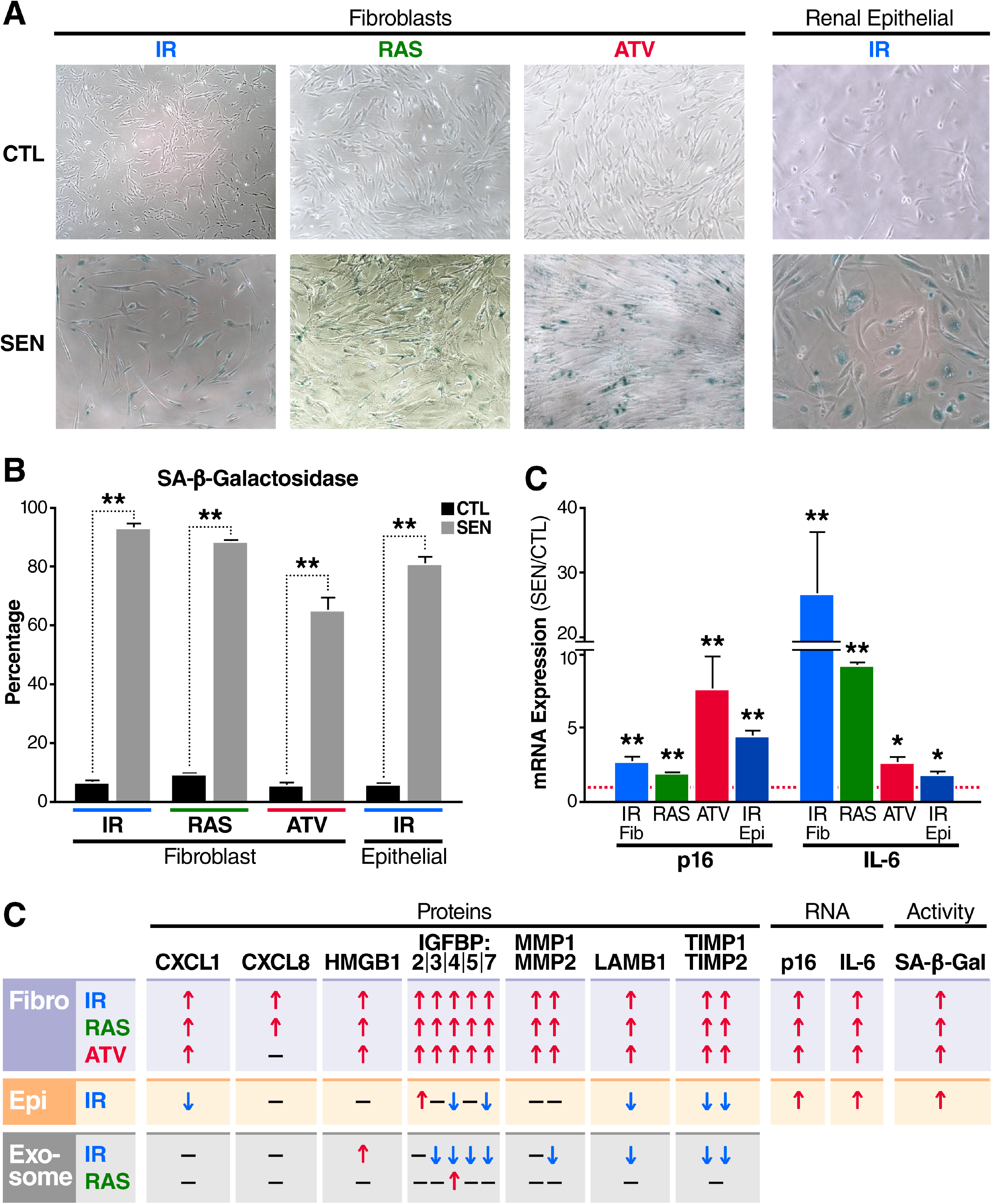
Comparison of proteomic and transcriptomic changes in the fibroblast SASP. Transcriptomic changes in the SASP of fibroblasts reported in a recent meta-analysis (23) (Hernandez-Segura et al., 2017) was compared with proteomic changes in the SASP of the current study. (A) Comparison of transcriptomic meta-analysis and proteomic analysis of secretomes in IR-induced senescent cells compared with non-senescent cells. (B) Venn diagram comparing RAS-induced senescence changes at the transcriptome and secreted proteome level. (C) Venn diagram of the core senescent transcriptome signature (genes changed at senescence regardless of inducer) versus changes common to IR and RAS induced senescence at the secreted proteome level. (D) Venn diagram comparing the senescent transcriptome and secreted proteome core signatures. IR = x-irradiation, RAS = RAS oncogene overexpression.

**Table S1:** Mass spectrometry quantification for each dataset as separate worksheets in a single excel workbook.

**Table S2:** Proteins with significantly increased secretion in response to all senescence-inducers.

**Table S3:** Proteins with significantly increased secretion in all cell types in response to all senescence inducers.

**Table S4:** Age-associated plasma proteins also present in the SASP as determined in our proteomics experiments.

**Table S5:** Reagents and Resources.

