## Supplementary material for "A Proteomic Atlas of Senescence-Associated Secretomes for Aging Biomarker Development": Table S5

### Supplemental Table 5 – Reagents and Resources

| REAGENT or RESOURCE | SOURCE | IDENTIFIER |
| --- | --- | --- |
| Chemicals, Peptides, and Recombinant Proteins | | |
| Atazanavir | MedChemExpress | HY-17367 |
| Pen-Strep | Gibco | 15070063 |
| DMEM | Gibco | 12430-054 |
| Trypsin-EDTA | Corning | 25-051-CI |
| Fetal Bovine Serum | Gibco | 2614079 |
| PBS | Gibco | 10010-23 |
| DMEM (phenol-red-free) | Gibco | 21063-029 |
| Acetonitrile (HPLC grade) | Burdick & Jackson | AH015 |
| Water (HPLC grade) | Burdick & Jackson | AH365 |
| Iodoacetamide | Sigma | I1149 |
| Dithiothreitol | Sigma | D9779 |
| Formic Acid | Sigma | 94318 |
| Triethylammonium bicarbonate | Sigma | T7408 |
| Urea | Thermo Scientific | 29700 |
| Trypsin (sequencing grade) | Promega | V5113 |
| HLB Oasis SPE cartridges | Waters | 186003908 |
| Amicon Ultra-15 Centrifugal Filter Units, 3kDa | MilliporeSigma | UFC900324 |
| Hyper Reaction Monitoring Peptide Standards | Biognosys | Kit-3003 |
| Sytox Green | Invitrogen | S7020 |
| DNase I Amp | Invitrogen | 18068015 |
| Critical Commercial Assays | | |
| Senescence-Associated-Beta-Galactosidase Staining Kit | BioVision | K320-250 |
| BCA Protein Assay Kit | Pierce | 23225 |
| MACSPlex Exosome Kit, human | Miltenyi | 130-108-813 |
| Lenti-X™ Tet-On® Advanced Inducible Expression System | Clontech | 632162 |
| PureLink Micro-to-Midi total RNA Purification System | Invitrogen | 12183018A |
| High-Capacity cDNA Reverse Transcription Kit | Applied Biosystems | 4368813 |
| Deposited Data | | |
| SASP Database | This paper | <www.SASPAtlas.com> |
| Proteomic SASP Panels | This paper, [Panorama](https://panoramaweb.org/project/home/begin.view?) (1) | <https://panoramaweb.org/project/Schilling/SASP_Atlas_Buck/begin.view?> |
| Proteomics data | This paper, [MassIVE](https://massive.ucsd.edu/ProteoSAFe/static/massive.jsp) | [MSV000083750](https://massive.ucsd.edu/ProteoSAFe/dataset.jsp?task=f0e0c2ef93c04521ba9656e07750a5fd) |
| Proteomics data | This paper, [ProteomeXchange](http://www.proteomexchange.org/) | [PXD013721](http://proteomecentral.proteomexchange.org/cgi/GetDataset?ID=PXD013721) |
| Pan-Human Spectral Library | Rosenberger et al., 2014 (2) |  |
| Human Proteome | [UniProt](https://www.uniprot.org/) | [UP00005640](https://www.uniprot.org/uniprot/?query=human&fil=reviewed%3Ayes+AND+organism%3A%22Homo+sapiens+%28Human%29+%5B9606%5D%22+AND+proteome%3Aup000005640&sort=score) |
| Plasma Aging Biomarkers | Tanaka et al., 2018 (3) |  |
| Gene Ontology | [The Gene Ontology Resource](http://geneontology.org/) |  |
| Human Kegg pathways | [KEGG PATHWAY Database](https://www.genome.jp/kegg/pathway.html) |  |
| Human WikiPathways | [WikiPathways](https://www.wikipathways.org/index.php/WikiPathways) |  |
| Human Reactome Pathways | [Reactome Pathway Database](https://reactome.org/) |  |
| Experimental Models: Cell Lines | | |
| IMR90 primary human lung fibroblasts | ATCC | CCL­-186 |
| human primary renal cortical epithelial cells (HRCE) | ATCC | PCS-400-011 |
| Software and Algorithms | | |
| Spectronaut X (version 12) | Biognosys, Rosenberger et al, 2014 (2) | <https://www.biognosys.com/shop/spectronaut-x> |
| Cytoscape (version 3.7.1) | Shannon et al., 2003 (4) | <https://cytoscape.org/> |
| ClueGO (version 2.5.3) | Bindea et al., 2009 (5) | <http://apps.cytoscape.org/apps/cluego> |
| R (version 3.5.2) | R Development Core Team, 2011 | <https://www.r-project.org>, RRID:SCR_001905 |
| Rstudio (version 1.0.136) | Rstudio: Integrated Development for R. RStudio, Boston, MA | <https://www.rstudio.com/>, RRID:SCR_000432 |
| VennDiagram (R) | Chen, 2018 (6) | <https://cran.r-project.org/web/packages/VennDiagram/index.html> |
| gplots (R) | Warnes et al., 2019 (7) | <https://cran.r-project.org/web/packages/gplots/index.html> |
| RColorBrewer (R) | Neuwirth, 2014 (8) | <https://cran.r-project.org/web/packages/RColorBrewer/index.html> |
| Scikit-learn (Python) | Pedregosa et al., 2011 (9) | <https://scikit-learn.org/stable/> |
| Other | | |
| SASP Atlas | This paper | <www.SASPAtlas.com> |
